# GPR183 targets lung-resident CD301b^+^ conventional dendritic cells type 2 to a subtissular TSLP – TSLP receptor mediated survival niche within the adventital cuff

**DOI:** 10.1101/2022.08.28.505379

**Authors:** L. Zhang, J. Yu, S. Spath, S. Sheoran, D Bejarano, M Vandestienne, AK Weier, T. Quast, M. Ibrahim, S Reimer, S Uderhardt, E. Mass, J. Hasenauer, A. Pfeifer, E. Kiermaier, W. Kolanus, S. Ziegler, A. Schlitzer

## Abstract

Conventional dendritic cells (cDCs) are strategically localized throughout non-lymphoid tissues. How such spatially regulated subtissular placement is achieved remains largely elusive. Here, we reveal that GPR183 targets CD301b^+^ cDC2 to a TSLP-dependent survival niche within the adventital cuff. We identified a close association of CD301b^+^ cDC2 with PDGFRα^+^ fibroblasts within the adventitial region of the lung. Genetic ablation of GPR183 within the cDC lineage leads to a selective loss of CD301b^+^ cDC2 in conjunction with the loss of CD301b^+^ cDC2 : fibroblast colocalization. Next bone marrow chimeric experiments and expression studies suggested adventitial fibroblasts as the main source of 7α,25 hydroxycholesterol, the natural ligand of GPR183. Single cell transcriptomic receptor ligand inference and subsequent genetic validation revealed TSLP receptor signalling as a crucial fibroblast derived CD301b^+^ cDC2 survival factor. These data expose a subtissular localization mechanism for tissue-specific functionalization of CD301b^+^ cDC2.

## Introduction

Conventional dendritic cells (cDCs) are the major antigen-presenting cells and crucial players for the induction of adaptive T-cell responses in mice and humans. cDCs can be separated into cDC1 and cDC2 according to their ontogeny, transcriptomic programs, function, and subtissular location (Ginhoux et al., 2009; Grajales-Reyes et al., 2015; Haniffa et al., 2012; Schlitzer et al., 2013; Schlitzer et al., 2015b). Functionally, cDC1 are specialized in the induction of immune responses against viruses and intracellular bacteria through the release of interleukin 12 (IL12) and the ability to cross-present antigen, whereas cDC2 release copious amounts of IL6, IL1b, and IL23 and induce immunity against extracellular bacteria and parasites (Everts et al., 2015; Haniffa *et al*., 2012; Schlitzer *et al*., 2013). Developmentally, cDC1 and 2 arise from transcriptionally specialized pre-cDC1 and 2 which terminally differentiate into cDC subsets under the influence of the inhabited tissue microenvironment (Cook et al., 2018; Grajales-Reyes *et al*., 2015; Schlitzer *et al*., 2015b; See et al., 2017). Along these lines, several molecules have been linked to cDC subset differentiation within peripheral tissues. Specifically, NOTCH2 (spleen, intestine) (Lewis et al., 2011; Satpathy et al., 2013), Lymphotoxin B (lung, intestine, dermis) (Kabashima et al., 2005; Satpathy *et al*., 2013) or GPR183 (spleen) (Baptista et al., 2019; Gatto et al., 2013; Li et al., 2016; Yi and Cyster, 2013) have been proposed to play crucial roles for the maintenance and/or development of cDC2 within the tissue microenvironment. In lung and skin functionally different cDC2 subsets have been defined according to their expression of the c-type lectin CD301b. CD301b^+^ cDC2 have been linked to the induction of Th2 immune responses, whereas the role of CD301b-cDC2 remains understudied (Kim et al., 2018; Kumamoto et al., 2009; Kumamoto et al., 2016; Kumamoto et al., 2013a; Murakami et al., 2013). However cellular components crucial to these subtissular niches remained largely elusive.

G protein-coupled receptors (GPCR) are sensors for extracellular cues, such as chemokines, cytokines, nutrients, or derivatives thereof in mononuclear phagocytes. However, candidates regulating subtissular placement of cDC subsets within their non-lymphoid organ tissue niches have not been identified (Gouwy et al., 2014). Among GPCRs, GPR183 is expressed on a variety of splenic resident cDC2, T, and B cells as well as on intestinal innate lymphoid cells, where it has been linked to a variety of different functions ranging from metabolism, effector function, differentiation, and immune cell migration (Baptista *et al*., 2019; Gatto et al., 2009; Gatto *et al*., 2013; Hannedouche et al., 2011; Jia et al., 2018; Kelly et al., 2011; Li *et al*., 2016; Wanke et al., 2017). Furthermore, GPR183 expression on various glia cells in the brain of mice undergoing neuroinflammation regulates disease outcome. GPR183 specifically recognizes 7α,25 hydroxycholesterol (7α,25-OHC). 7α,25-OHC is synthesized from cholesterol via an enzymatic cascade involving the rate-limiting enzymes 25-hydrolase (CH25H) and the P450, family 7, subfamily b, polypeptide 1 (CYP7B1) protein. Degradation of 7α,25-OHC is mediated by the protein hydroxy-delta-5-steroid dehydrogenase, 3 beta- and steroid delta-isomerase 7 (HSD3B7), which degrades 7α,25-OHC into GPR183 non-reactive bile acid precursors. The lung is a cholesterol-rich environment and as *Ch25h* is abundantly expressed within alveolar macrophages (Gautier et al., 2012; Jia *et al*., 2018; Madenspacher et al., 2020), it is plausible that the 7α,25-OHC : GPR183 signalling axis plays an important role in the subtissular guidance of cDC subsets towards their respective microenvironmental niches within the lung.

Further evidence for an subtissular guidance role of GPR183 stems from its role in the positioning of T and B cells within the spleen and the lymph node, where it is crucial for correct positioning of T follicular helper cells around the follicle and for entry of B-cells into the germinal center (Gatto *et al*., 2009; Hannedouche *et al*., 2011; Kelly *et al*., 2011; Li *et al*., 2016). Additionally, small intestinal innate lymphocytes depend on GPR183 signalling for optimal functional positioning within intestinal lymphoid follicles (Emgård et al., 2018b). Therefore, it is compelling to investigate the role of GPR183 as a central subtissular guidance factor for the spatial separation of cDC subsets in cholesterol-rich organs, such as the lung.

Here, using a cDC-specific knockout mouse model for *Gpr183*, high dimensional spatial analysis, and transcriptomic ligand : receptor interaction prediction, we define 7α,25-OHC and its receptor GPR183 as an subtissular guidance factor for lung-resident CD301b^+^ cDC2. Loss of *Gpr183* on cDCs or of *Ch25h* in non-immune cells leads to loss of CD301b^+^ cDC2 within the lung. cDC2 loss could not be attributed to defective DCpoesis or to a direct role of GPR183 in the maintenance of CD301b^+^ cDC2s or their progenitors. cDC-specific GPR183 deficiency leads to increased CD301b^+^ cDC2 apoptosis and abrogation of *in situ* CD301b^+^ cDC2 proliferation. Spatial analysis of murine homeostatic lungs identified that CD301b^+^ cDC2 expressed GPR183 and its stromal-derived ligand 7α,25-OHC as a guidance signal for CD301b^+^ cDC2 into a PDGFRα^+^Sca1^+^ fibroblastic niche. Receptor-ligand prediction of the PDGFRα^+^ fibroblast: CD301b^+^ cDC2 circuit coupled to genetic validation revealed that thymic stromal lymphopoietin (TSLP) and its receptor act as a fibroblast-derived subtissular CD301b^+^ cDC2 survival signal to which GPR183-mediated attraction acts as an subtissular navigation beacon.

## Results

### CD301b^+^ cDC2 enrich in adventitial areas of the murine lung

To understand the subtissular distribution of tissue-resident cDC subsets within the lung microenvironment, we investigated the distribution of cDC1, and CD301b^+^ and CD301b^-^ cDC2 in the homeostatic murine lung using co-detection by indexing (CODEX), which enabled high dimensional imaging of fresh frozen lung sections (Figure 1A, B, **Figure S1A, B)**. This analysis showed that CD301b^+^ cDC2 enrich within adventitial cuffs as compared to other anatomical areas, such as bronchi or alveolar areas (Figure 1C), whereas cDC1 and CD301b^-^ cDC2 were evenly distributed across the anatomical areas investigated.

**Figure 1.**
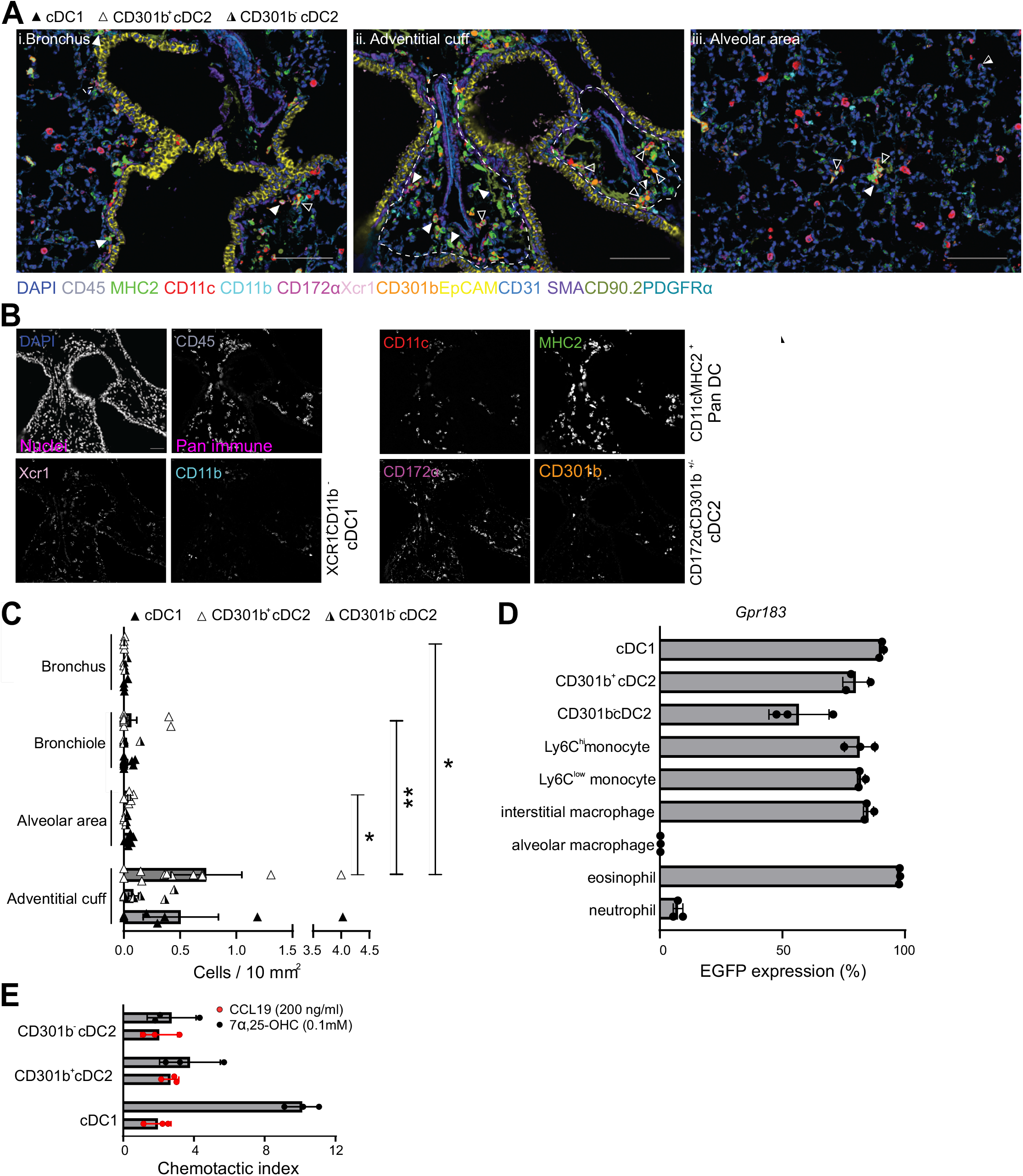
cDC1 and cDC2 localize in distinct microanatomical areas of the murine lung. (A) Representative CODEX images of bronchial (i), adventitial (ii), and alveolar areas (iii). Adventitial cuff (delineated by dashed lines in (ii)) were separated from bronchial areas using EpCAM and PDGFRα signals as a reference. Solid arrows indicate cDC1, empty arrows indicate CD301b^+^ cDC2, and half empty arrows indicate CD301b^-^ cDC2. Scale bars represent 200 µm. (B) Representative images of the single stainings of cDC related markers used for CODEX. Scale bars represent 100µm. (C) Quantification of cDC1, CD301b^+^ and CD301b^-^ cDCs in discrete anatomical regions. Absolute cell numbers of cDC1 (solid arrows), CD301b^+^ cDC2 (empty arrows), and CD301b^-^ cDC2 (half empty arrows) in region were calculated and normalized by the area. Each dot represents the normalized value of each annotated region. (D) GPR183 expression on myeloid subsets as assessed by EGFP expression in *Gpr183*^flox-EGFP^ mice; n = 3. (E) Transwell migration of sorted pulmonary cDC1, CD301b+ and Cd301b-cDC2 in WT mice to CCL19 and 7α,25-OHC, n = 3. Each dot refers to one mouse in (D) and (E). Data are represented as means ± SD. ∗p < 0.05, ∗∗p < 0.01 was assessed by two-tailed non-paired Mann-Whitney test.

These results prompted us to investigate how CD301b^+^ cDC2 positioning is regulated within the lung microenvironment. Therefore, we investigated whether myeloid cells including cDC1, CD301b^+^, and CD301b^-^ cDC2 express GPR183, within the cholesterol-rich environment of the lung using a GPR183 reporter mouse model (Andersson et al., 2017). This analysis showed that both cDC1, CD301b^+^ and CD301b^-^ cDC2 abundantly expressed GPR183 (Figure 1D**, Figure S1C, D)**. Next, we wanted to investigate whether GPR183 expression leads to functional migration of *ex vivo* isolated cDC1, CD301b^+^ and CD301b^-^ cDC2 towards the the natural GPR183 ligand 7α,25-OHC as previously reported *in vitro* (Gatto *et al*., 2013). To study this we purified lung resident cDCs, seeded them on transwell filter, and incubated them in the presence or absence of 7α,25-OHC in the lower chamber or with CCL19 as a positive control. After the culture period, we examined the amount of cDC1, CD301b^+^, and CD301b^-^ cDC2 within the lower chamber using flow cytometry. This analysis revealed that all three lung-resident cDC subsets (cDC1, CD301b^+^, and CD301b^-^ cDC2) migrated efficiently in response to 7α,25-OHC (Figure 1E) indicating GPR183 as a possible subtissular guidance cue for cDC subsets in the lung.

### cDC-specific deletion of *Gpr183* leads to a reduction of lung resident CD301b^+^ cDC2

To test this hypothesis *in vivo* we generated a cDC lineage-specific *Gpr183* deficient mouse model (*Zbtb46*^cre^*Gpr183*^flox-EGFP^) and examined the cellular composition of the homeostatic lung and spleen by flow cytometry (Figure 2A-C**, Figure S2A-D)**. In the lung, we observed a specific reduction of lung-resident CD301b^+^ and CD301b^-^ cDC2, whereas other mononuclear phagocyte subsets, including cDC1, remained unaffected (Figure 2A-C**, Figure S2A, B).** Additionally we found a specific reduction of splenic Esam^-^ and Esam^+^ cDC2 in *Zbtb46*^cre^*Gpr183*^flox-EGFP^ mice as previously published ((Lu et al., 2017b)**, Figure S2C, D)** as previously published. Next, we assessed the cDC2 content of precision-cut lung slices generated from wildtype and *Gpr183*^-/-^ mice using confocal microscopy. This analysis confirmed a reduction in overall numbers of cDC2s (CD172a^+^CD11c^+^CD88^-^) in the lung (Figure 2D, E). To further characterize the nature of the observed decrease in lung-resident CD301b^+^ and CD301b^-^ cDC2 in *Zbtb46*^cre^*Gpr183*^flox-EGFP^ mice and to rule out off-target effects of the *Zbtb46*^cre^ mouse model (Meredith et al., 2012; Satpathy et al., 2012), we transferred bone marrow isolated from CD45.2^+^*Zbtb46*^cre^*Gpr183*^flox-EGFP^ mice into lethally irradiated CD45.1^+^ wild-type mice and analyzed the lungs of those recipients 12 weeks after bone marrow transfer. This analysis showed a specific reduction of lung resident CD301b^+^ cDC2 in chimeras transplanted with CD45.2^+^*Zbtb46*^cre^*Gpr183*^flox-EGFP^ bone marrow but not in control chimeras (Figure 2F, G). These findings clearly establish *Gpr183* as a cDC intrinsic regulator of lung resident cDC2 homeostasis.

**Figure 2.**
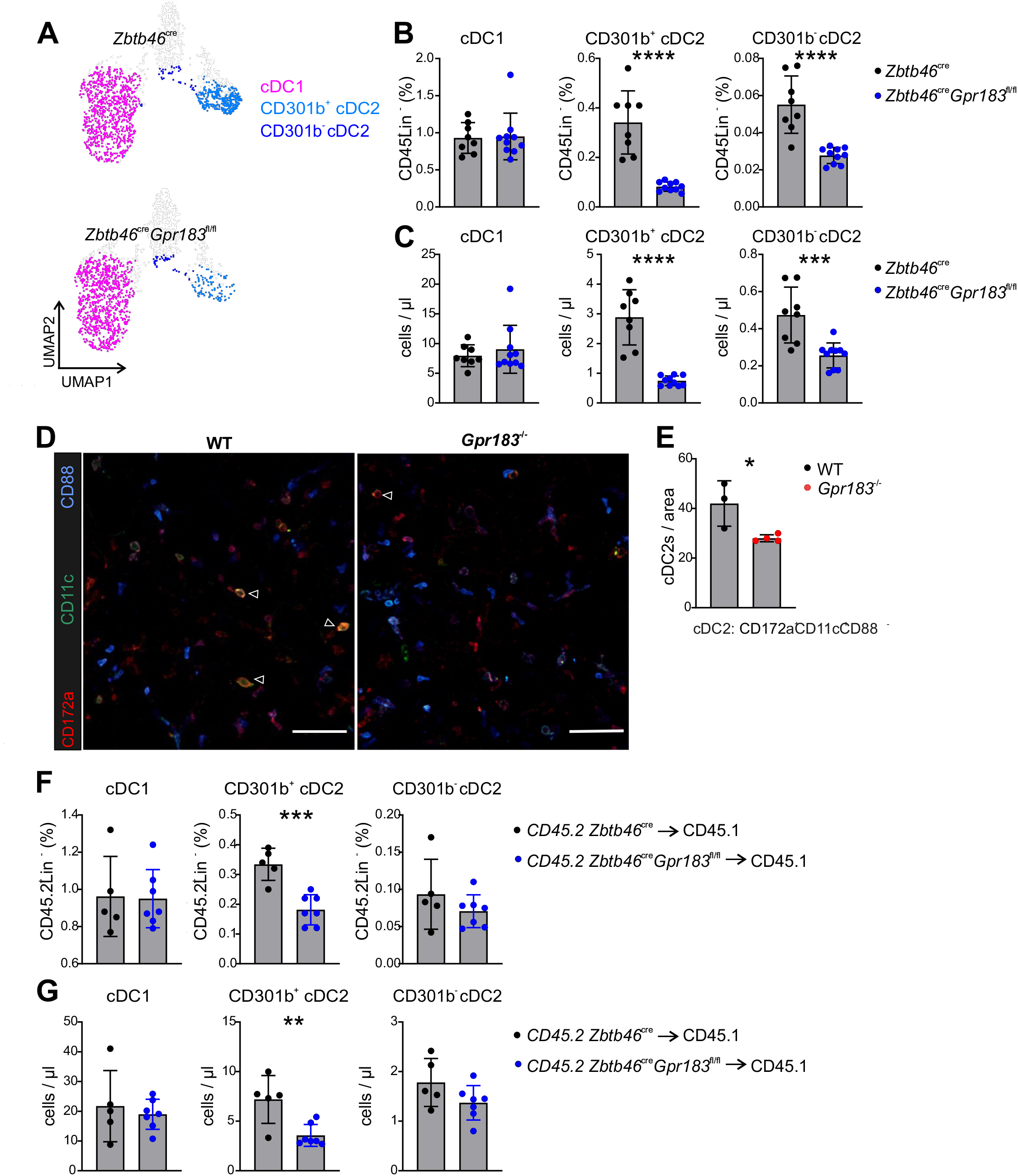
cDC lineage specific deletion of GPR183 leads to a reduction in lung resident cDC2. (A) UMAP visualization of pulmonary CD45^+^Lin^-^CD64^-^Ly6C^-^MHC2^+^CD11c^+^CD24^+^ cells from *Zbtb46*^cre^ and *Zbtb46*^cre^*Gpr183*^flox-EGFP^ mice. Colour indicates classification based on the expression of canonical markers. FACS analysis of cDC1, CD301b^-^ and CD301b^+^ cDC2 in lungs of *Zbtb46*^cre^ (black) and *Zbtb46*^cre^*Gpr183*^flox-EGFP^ (blue) mice. Plots show the frequency (B) and cell concentration (C) of cDC1, CD301b^-^ and CD301b^+^ cDC2; n = 8 - 10. (D) Representative confocal images of precision cut lung slices of WT and *Gpr183*^-/-^ mice. Slices were immunostained using anti-CD11c (green), anti-CD172a (red), and anti-CD88 (blue) antibodies to visualize cDC2. Empty arrows indicate cDC2s. Scale bars represent 400 μm. (E) Quantification of cDC2 number per area (531μm x 531μm) in lung slices from WT (black dots) and *Gpr183*^-/-^ (red dots) mice; n = 3 or 4. FACS analysis of cDC1, CD301b^-^ and CD301b^+^ cDC2 in lungs of CD45.1 bone marrow recipients transplanted with either *Zbtb46*^cre^ (black) or and *Zbtb46*^cre^*Gpr183*^flox-EGFP^ (blue) bone marrow. Plots show the frequency (F) and cell concentration (G) of cDC1 and cDC2 subsets; n = 5 - 7. Each dot refers to one mouse. Data in (B) and (C) are pooled from two independent experiments. Data in (F) and (G) are representative data of two independent experiments. Data are represented as means ± SD. ∗p < 0.05, ∗∗p < 0.01, ∗∗∗p < 0.001, ∗∗∗∗p < 0.0001 by unpaired T-test.

### DC lineage specific ablation of *Gpr183* does not affect cDC development

Tissue resident cDC subsets are continuously replenished by pre-cDC1 and 2, which in turn are produced from CDPs within the bone marrow (BM) (Grajales-Reyes *et al*., 2015; Schlitzer et al., 2015a). In order to dissect the mechanism(s) involved in the observed reduction of lung-resident cDC2 subsets in the absence of *Gpr183,* we first analysed the expression of *Gpr183* within the BM, blood and lung pre-cDC compartment using *Gpr183*^flox-EGFP^ reporter mice (Figure 3A**, Figure S3A-C)**. This analysis showed expression of *Gpr183* on 45.13 ± 4.27 % of pre-cDC1 and 91.65 ± 0.94 % pre-cDC2 in the bone marrow, 93.40 ± 0.29 % of pre-cDC1 and 98.46 ± 0.15 % pre-cDC2 in the blood and 71.87 ± 1.27 % of pre-cDC1 and 88.10 ± 0.94 % pre-cDC2 in the lung (Figure 3A), suggesting that GPR183 and its ligand 7α,25-OHC directly act on pre-cDCs either in the lung or the periphery. To test this we quantified MDP, CDP, pre-cDC 1 and 2 in the BM, blood and lung of *Zbtb46*^cre^*Gpr183*^flox-EGFP^ ^or^ WT mice using flow cytometry. Here no differences in abundance of MDP, CDP, pre-cDC 1 and 2 in the BM, blood or lung between control and *Gpr183*^-/-^ mice was found (**Figure S3D, E).** The above analysis ruled out a direct effect of GPR183 on the recruitment of pre-cDC subsets towards the lung, however did not directly address its effect on DC development in the bone marrow. To investigate this we generated DCs from hematopoietic stem cells in the presence of FLT3 ligand. Cells were treated with either vehicle, 7α,25-OHC or NIBR189 (GPR183 antagonist) (Hannedouche *et al*., 2011) on day 3 and 5 of the 7-day culture period. On day 7 we utilised flow cytometric analysis and investigated the abundance of cDC1 and cDC2 subsets within our *in vitro* culture. Here no differences were found in the *in vitro* generation of cDC subsets when treated with either of the regimes mentioned above (**Figure S3F, G)**, thereby showing that GPR183 signalling does not affect cDC development *in vitro* or *in vivo*.

**Figure 3.**
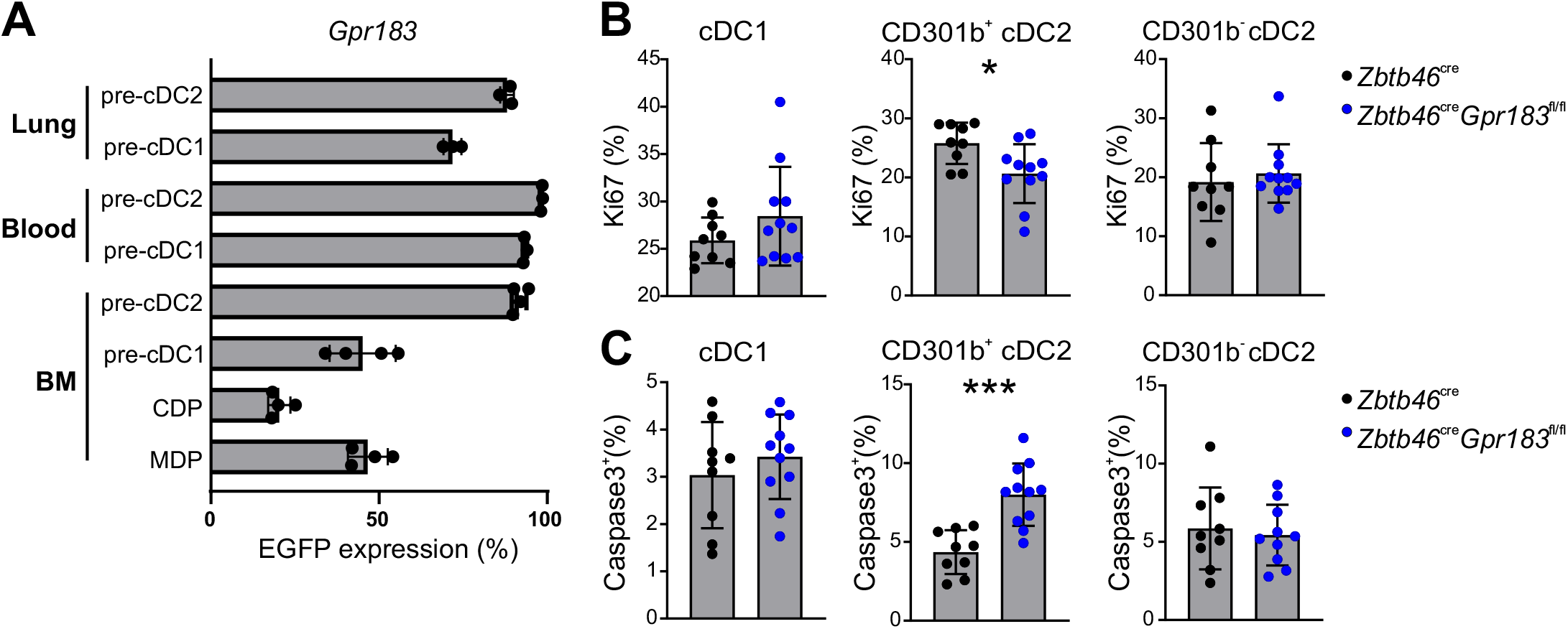
DC specific loss of GPR183 abolishes *in situ* proliferation of CD301b^+^ cDC2 and induces apoptosis. (A) Frequency of *Gpr183* expressing MDP, CDP, pre-cDC, pre-cDC1 and pre-cDC2 in BM, blood and lung of *Gpr183*^flox-EGFP^ mice; n = 3 or 4. (B) Frequency of Ki67^+^ ccDC1, CD301b^+^ and CD301b^−^ cDC2 in lungs of *Zbtb46*^cre^*Gpr183*^flox-EGFP^ (blue) and *Zbtb46*^cre^ (black) mice; n = 9 - 11. (C) Frequency of active caspase 3^+^ cDC1, CD301b^+^ cDC2 and CD301b^−^ cDC2 in lungs of *Zbtb46*^cre^*Gpr183*^flox-EGFP^ (blue) and *Zbtb46*^cre^ (black) mice; n = 9 or 11. Each dot refers to one mouse. Data in (A) are representative data of two independent experiments. Data in (B) and (C) are pooled from two independent experiments. Data are represented as means ± SD. ∗p < 0.05, ∗∗∗p < 0.001 by unpaired T-test.

### DC specific loss of GPR183 abolishes proliferation of CD301b^+^ cDC2 and induces apoptosis *in situ*

As loss of GRP183 on pre-cDCs did not affect recruitment to the lung or development of cDCs we assessed proliferation and apoptosis of pulmonary pre-cDCs and cDCs using flow cytometry. Here loss of GPR183 did not affect proliferation of pulmonal pre-cDC1, 2, cDC1 and CD301b^-^ cDC2. Vice versa we observed a decrease in proliferative capacity of CD301b^+^ cDC2 as assessed by expression of Ki67 (**Figure S3H,** Figure 3B). Next to understand whether the reduction of CD301b^+^ cDC2 proliferation within the lung coincides with increased apoptosis of remaining CD301b^+^ cDC2s in *Zbtb46*^cre^*Gpr183*^flox-EGFP^ we assessed abundance of activated Caspase 3 as a proxy for apoptosis of pulmonal pre-cDC1, 2, cDC1 CD301b^+^ cDC2 and CD301b^-^ cDC2. Here no effect on activation of Caspase 3 was observed in pre-cDC1, 2, cDC1 and CD301b^-^ cDC2 isolated from *Zbtb46*^cre^*Gpr183*^flox-EGFP^ as compared to WT mice (**Figure S3I,** Figure 3C). However assessing activation of Caspase 3 in CD301b^+^ cDC2 isolated from *Zbtb46*^cre^*Gpr183*^flox-EGFP^ showed an increase in activated Caspase 3^+^ cells compared to WT CD301b^+^ cDC2 (Figure 3C). Taken together these data reveal that loss of GPR183 on CD301b^+^ cDC2 affects their ability of *in situ* maintenance by decreasing cellular proliferation and increasing apoptosis independent of direct GPR183 – ligand interaction.

### 7α,25-OHC production by radioresistant non-immune cells maintains lung resident cDC2 compartment

Previous reports indicated splenic and intestinal non immune cells as important sources of 7α,25-OHC. However within the lung the source of 7α,25-OHC remained unknown (Chu et al., 2018; Emgård *et al*., 2018b; Lu *et al*., 2017b; Rodda et al., 2018). Thus we investigated expression of *Ch25h*, *Cyp7b1* as producing and *Hsd3b7* as degrading enzymes of 7α,25-OHC respectively, to further understand how the GPR183 : 7α,25-OHC – CH25H signalling circuit guards CD301b^+^ cDC2 maintenance and possibly positioning in the murine lung (Figure 4A**, Figure S4A)**. *Realtime* PCR (RT-PCR) analysis showed that *Ch25h* and *Hsd3b1* were expressed in all investigated cell types except epithelial cells, whereas Cyp7b1 was specifically expressed by PDFGRα^+^SCA1^+^ fibroblasts (Figure 4A), making it likely that fibroblasts are producing 7α,25-OHC in the lung microenvironment, in line with previously reported findings in the intestine, spleen and lymph node (Emgård et al., 2018a; Emgård *et al*., 2018b; Gatto *et al*., 2013; Hannedouche *et al*., 2011). To validate these findings *in vivo* we utilized *Ch25h* deficient mice and analysed the abundance of pulmonary cDC1, CD301b^+^ and CD301b^-^ cDC2 by flow cytometry. Here we found a reduction in frequency and absolute number of CD301b^+^ and CD301b^-^ cDC2s in *Ch25h*^-/-^ mice as compared to WT counterparts, whereas cDC1 frequency and number remained unaffected (Figure 4B, C, D). In order to further pinpoint the cellular source of 7α,25-OHC, we transferred CD45.1^+^ BM cells into lethally irradiated *Ch25h*^-/-^ recipient mice and analysed the abundance of cDC1, CD301b^+^ and CD301b^-^ cDC2s 12 weeks after BM transplantation using flow cytometry. Here lung resident CD301b^+^ and CD301b^-^ cDC2s but not cDC1 were significantly reduced in mice with a Ch25h^-/-^ non-immune compartment (Figure 4E, F). Next to understand the role of immune cells for the production of 7α,25-OHC in the lung we transferred *Ch25h*^-/-^ bone marrow into lethally irradiated CD45.1^+^ recipients and analysed abundance of lung resident cDC subsets by flow cytometry. This analysis revealed that *Ch25h* deficiency within the immune cell compartment had no effect on frequencies or absolute cell numbers of pulmonal cDC subsets (**Figure S4B, S4C)**. These results suggested 7α,25-OHC production by stromal cells, likely adventitial fibroblasts, is critical for lung-resident CD301b^+^ cDC2 maintenance within the lung microenvironment.

**Figure 4.**
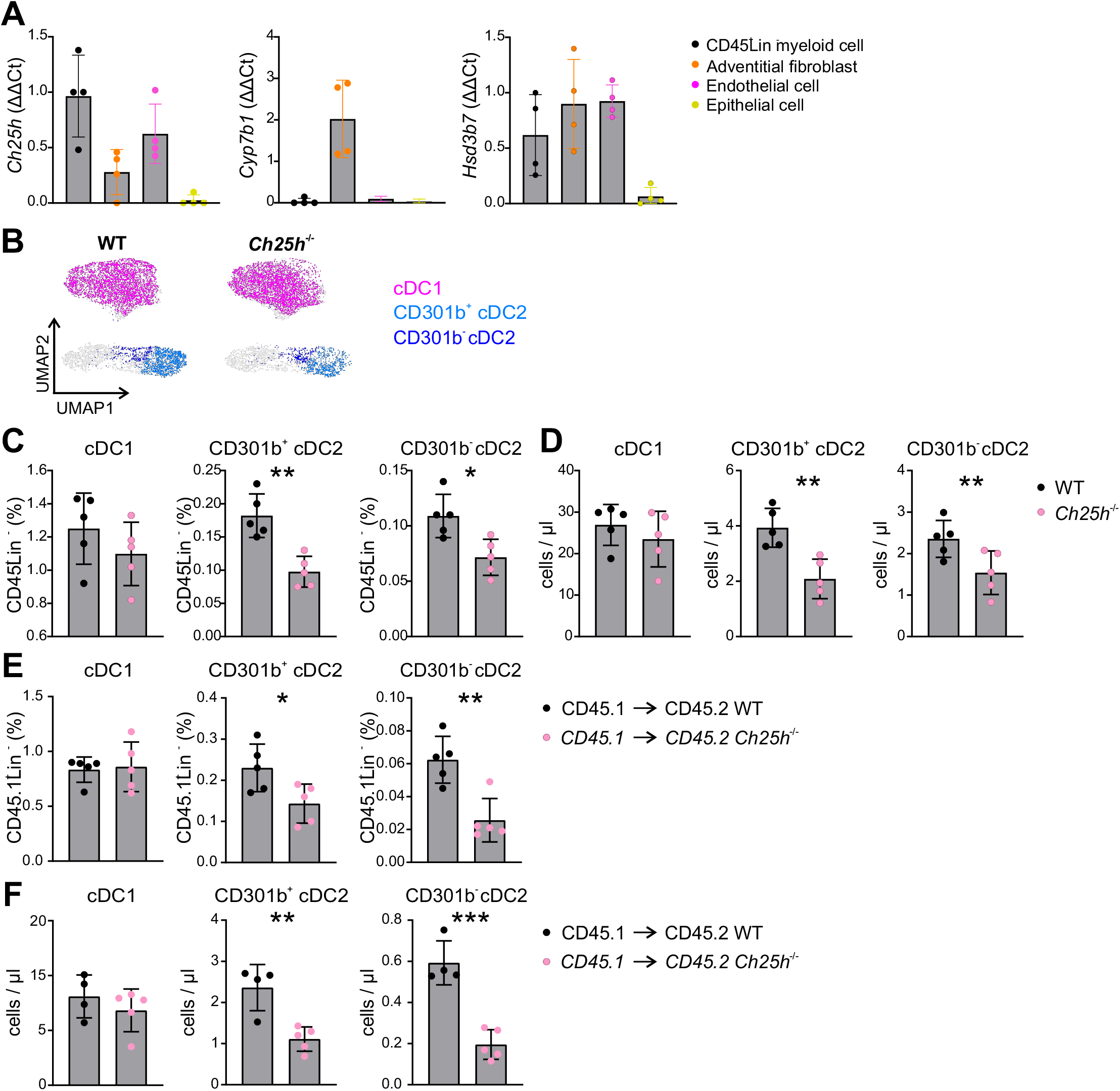
CH25H dependent production of 7α,25-OHC by radioresistant non-immune cells maintains lung resident cDC2 compartment. (A) RT-PCR analysis of *Ch25h*, *Cyp7b1* and *Hsd3b7* transcript abundances in purified stromal cell subsets and CD45^+^Lin^-^ cells from murine WT lungs. Plots show relative quantification normalized to the expression of the housekeeping gene *PPIA*; n = 4. (B) UMAP visualization of pulmonary CD45^+^Lin^-^CD64^-^Ly6C^-^MHC2^+^CD11C^+^CD24^+^ cells from WT and *Ch25h*^-/-^ mice. Colour indicates classification based on expression of canonical markers. FACS analysis of cDC1, CD301b^-^ and CD301b^+^ cDC2 in lungs of *Ch25h*^-/-^ (pink) and WT (black) mice. Plots show the frequency (C) and cell concentration (D) of cDC1, CD301b^-^ and CD301b^+^ cDC2 subsets; n = 5. FACS analysis of cDC1, CD301b^-^ and CD301b^+^ cDC2 in lungs of CD45.1^+^ bone marrow recipients transplanted with either CD45.2^+^*Ch25h*^-/-^ (pink) or WT (black) donor bone marrow. Plots show the frequency (E) and cell concentration (F) of cDC1, CD301b^-^ and CD301b^+^ cDC2; n = 4 - 5. Each dot refers to one mouse. Data are represented as means ± SD. ∗p < 0.05, ∗∗p < 0.01, ∗∗∗p < 0.001 by unpaired T-test.

### GPR183 guides pulmonary CD301b^+^ cDC2 into a adventitial fibroblastic pro-survival niche

To test the hypothesis that *Cyp7b1^+^Ch25h^+^* PDFGRα^+^SCA1^+^ fibroblasts attract lung-resident CD301b^+^ cDC2s and support their survival within a specific subtissular niche, we purified c CD301b^+^ and CD301b^-^ cDCs from WT lungs and studied migration in a co-culture with WT or *Ch25h*^-/-^ lung PDFGRα^+^SCA1^+^ fibroblasts *ex vivo*. This experiment revealed that WT but not *Ch25h*^-/-^ PDFGRα^+^SCA1^+^ fibroblasts attracted CD301b^+^ and CD301b^-^ cDC2 (Figure 5A). PDFGRα^+^SCA1^+^ lung fibroblasts are primarily found within adventitial cuffs (Dahlgren et al., 2019). Prior, adventitial cuffs have been described as immune cell hotspots within the lung microenvironment (Cautivo et al., 2020; Dahlgren *et al*., 2019). To assess whether CD301b^+^ cDC2 are localized close to fibroblasts *in vivo* we utilized whole mount live section confocal microscopy analysis visualizing cDC2 in conjunction with PDGFRα^+^ fibroblasts. This revealed that in WT lungs cDC2 were found in close proximity to PDFGRα^+^ fibroblasts. Next we investigated if loss of GPR183 affects co-localization of cDC2 and fibroblasts. Here we found that ablation of Gpr183^-/-^ only affected cDC2 which were in close proximity to PDGFRα^+^ fibroblasts but not cDC2 which were further away from fibroblasts (Figure 5B, C). Collectively these data suggest a GPR183 dependent CD301b^+^ cDC2 : PDFGRα^+^SCA1^+^ fibroblast interaction within the lung, crucial to the survival of CD301b^+^ cDC2.

**Figure 5.**
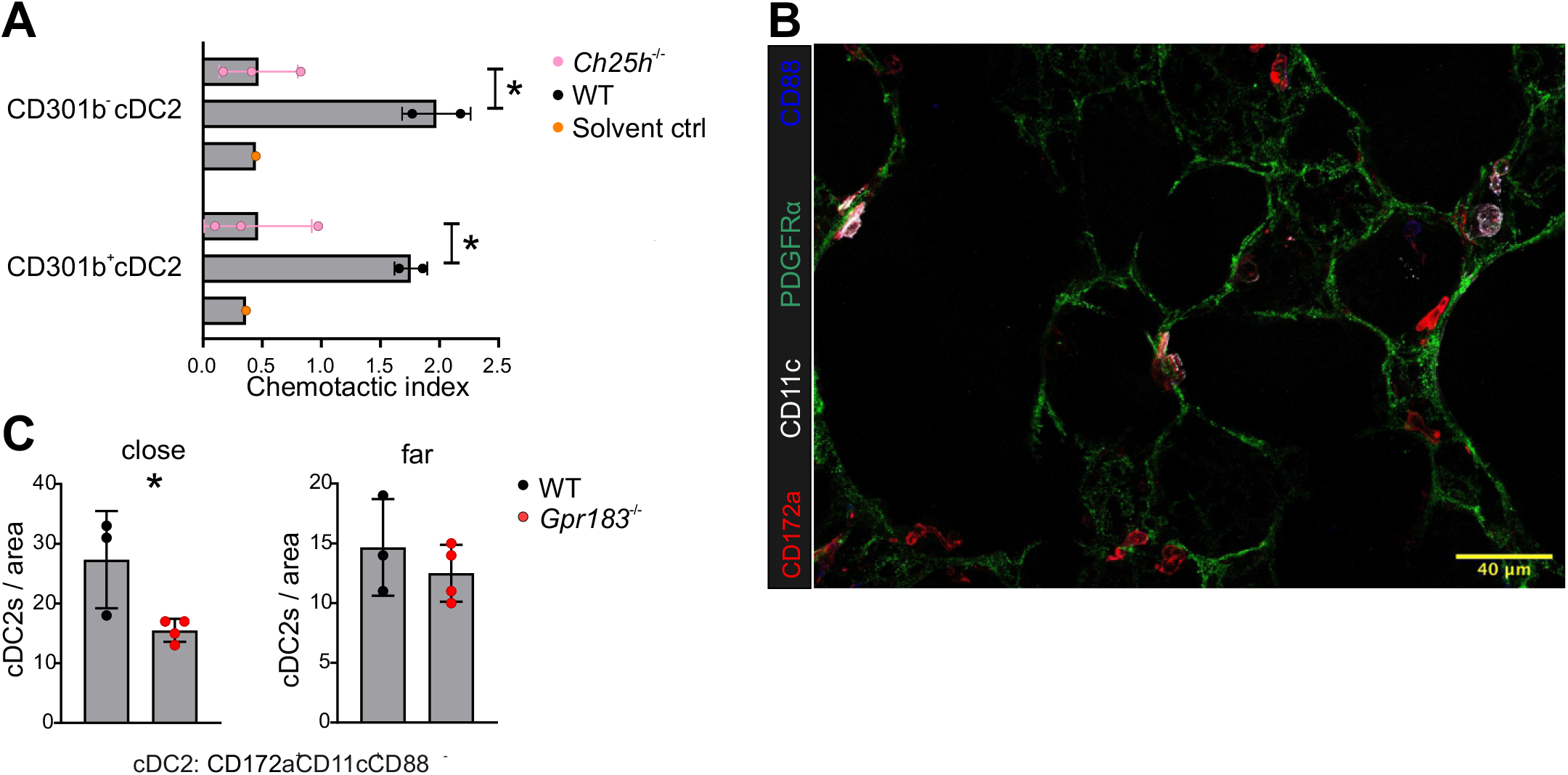
GPR183 guides pulmonary CD301b^+^ cDC2 into a fibroblastic pro-survival niche. (A) *In vitro* migration of purified WT pulmonal CD301b^-^ and CD301b^+^ cDC2 towards PDGFRα^+^ fibroblasts purified from WT (grey, n = 2) or *Ch25h*^-/-^ (pink, n = 3) mice, black bar represents solvent control (n = 1). (B) Representative confocal images of precision cut lung slices of WT mice. Slices were immunostained using anti-CD11c (white), anti-CD172a (red), anti-PDGFRα (green), and anti-CD88 (blue) antibodies to visualize cDC2s and fibroblasts. Scale bar represents 40 μm. (C) Quantification of the number of cDC2 which are in close and far distance to fibroblasts in precision cut lung slices; n = 3 or 4. Each dot refers to one mouse. Data are pooled from two independent experiments. Data are represented as means ± SD. ∗p < 0.05 by unpaired T-test.

### CD301b^+^ cDC2 : adventitial fibroblast receptor – Ligand interaction prediction reveals TSLP as crucial fibroblast derived survival factor for pulmonary CD301b^+^ cDC2

Our previous data indicated that direct GPR183 - 7α,25-OHC signalling does not mediate a direct pro-survival or developmental effect on lung-resident CD301b^+^ cDC2 *in vivo* and *in vitro* but directs CD301b^+^ cDC2s to an adventitial fibroblast niche, which further supports their survival. To understand the nature of this adventitial fibroblast : CD301b^+^ cDC2 interaction we investigated a previously published dataset comprised of 13823 single cells isolated from two healthy lungs of *Col1a1*-GFP reporter mice (Tsukui et al., 2020). *Col1a1* expression in the lung marks all fibroblasts and smooth muscle cells first identified as *GFP*^+^*Col1a1*^+^ cells from the data (Figure 6A). This identified 5 clusters corresponding to *GFP*^+^*Col1a1*^+^ fibroblasts (cluster 0, 1, 3, 6, 8) (Figure 6A**, left panel)** and cluster 19 as *Acta2*^+^*Myh11*^+^ smooth muscle cells (**Figure S5A)**. Our and other previous data (Dahlgren *et al*., 2019) showed that cDC2 closely align with PDGFRα^+^ fibroblasts in the lung (Figure 1A, 5C), hence we curated *GFP*^+^*Col1a1*^+^ lung fibroblast clusters using *Pdgfrα* expression and identified clusters 0-5, 8 and 10 as *Pdgfrα*^+^ lung fibroblasts **(Figure S5B)**. Next to identify adventitial fibroblasts we asked which *Pdgfrα*^+^ fibroblast sub-clusters express *Pi16* and *Gli1*, two established markers of adventitial lung fibroblasts (Dahlgren *et al*., 2019; Tsukui *et al*., 2020). This analysis revealed that clusters 3, 4 and 5 were adventitial fibroblasts (Figure 6B**, S5C)**. CH25H and CYP7B1 are key enzymes for the production of 7α,25-OHC. Therefore, we explored expression of those molecules within adventitial fibroblast clusters 3, 4 and 5. This showed co-expression of *Ch25h* and *Cyp7b1* within adventitial fibroblast clusters 3 and 5 (Figure 6C**, S5D)**. Next we identified *CD301b*^+^*Gpr183*^+^ cDC2 (in cluster 15; Figure 6D, **S5E)** from the same dataset using expression of the cDC-specific markers (*Itgae*, *Cd86*, *Mgl2* and *Gpr183*) and predicted interactions between *CD301b*^+^*Gpr183*^+^ cDC2 and cluster 3 and 5 respectively using NicheNet (Browaeys et al., 2020). This analysis identified various specific and shared (grey) interactions, between adventitial fibroblast clusters 3 (magenta), 5 (green) and *CD301b*^+^*Gpr183*^+^ cDC2 respectively, such as *ApoE* and *Lrp1* for cluster 3 or *Il6* and *Il6ra* for cluster 5 (Figure 6E). Next we ask whether shared interactions (grey) between cluster 3 and 5 exist, as both clusters expressed *Cyp7b1* and thus likely were both responsible for the production of 7α,25-OHC. This analysis revealed several shared interactions of clusters 3, 5 and *CD301b*^+^*Gpr183*^+^ cDC2. Among those the *Tslp* : *Crlf2* interaction had the highest prior interaction potential value and showed robust expression on both cluster 3 and 5 (Figure 6E), which was also validated in a second single cell transcriptomics dataset of murine the lung (Raredon et al., 2019) **(Figure S5F, S5G)**.

**Figure 6.**
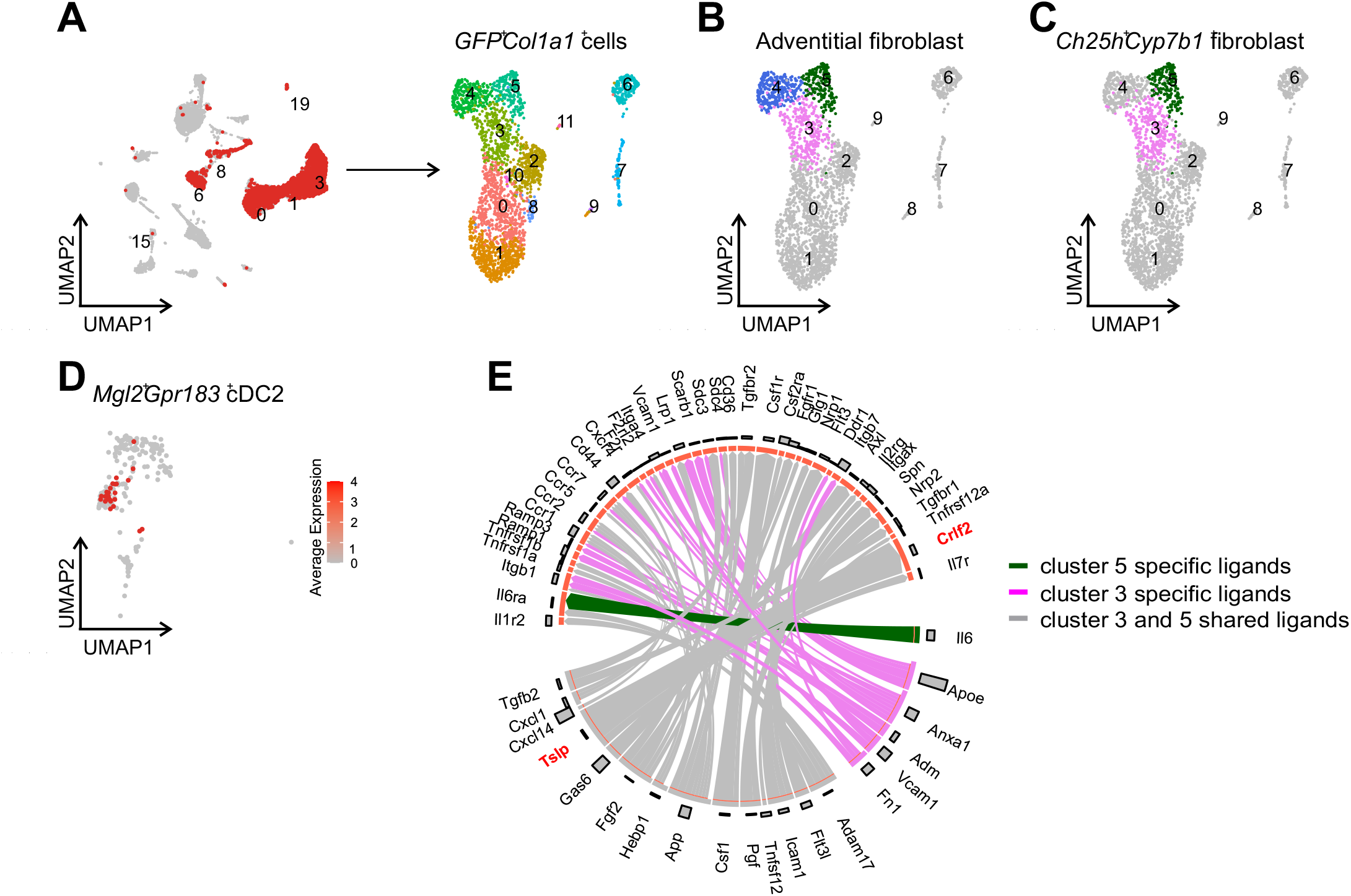
Receptor – Ligand interaction prediction reveals TSLP as crucial fibroblast derived survival factor for pulmonary cDC2. (A) UMAP embedding of 13823 cells from untreated murine lungs (GSE132771). Cluster 15 is annotated as cDCs (left panel). *GFP*^+^*Col1a1*^+^ cells are coloured in red (left panel) and further re-clustered into 12 clusters (right panel). (B) Identification of 3 sub-clusters of adventitial fibroblasts in *GFP*^+^*Col1a1*^+^ cells. (C) UMAP embedding of 2 *Ch25h*^+^*Cyp7b1*^+^ adventitial fibroblast sub-clusters within *GFP*^+^*Col1a1*^+^ cells. (D) Identification and UMAP embedding of *Mgl2*^+^*Gpr183*^+^ cDC2 coloured in red within cluster 15. (E) NicheNet analysis of potential interactions between receptors expressed on *Mgl2*^+^*Gpr183*^+^ cDC2 and ligands expressed on adventitial fibroblast from cluster 3 (violet), 5 (green) or from both (grey). Expression of ligands and receptors corresponds to bars represented in (E)

### DC specific ablation of TSLP receptor leads to a reduction in pulmonary CD301b^+^ cDC2

In order to experimentally validate the prediction that fibroblastic TSLP is important for CD301b^+^ cDC2 maintenance we first confirmed protein level expression of TSLP receptor on CD301b^+^ cDC2 and of TSLP released by lung fibroblasts using flow cytometry and ELISA, respectively (Figure 7A, B). This analysis showed that TSLP receptor was abundantly expressed on both CD301b^+^ and CD301b^-^ cDC2 during homeostasis in the lung (Figure 7A). Secondly, notable levels of secreted TSLP could be measured in homogenates of WT lungs and supernatants of fibroblasts from murine WT lungs after seven days of *ex vivo* culture (Figure 7B). These results prompted us to genetically validate the role of adventitial fibroblast derived TSLP for lung CD301b^+^ cDC2 homeostasis. Thus we generated a DC lineage specific deletion of the TSLP receptor (*CD11c*^cre^*Crlf2*^flox^) and analysed lungs of control and DC-specific TSLP receptor deleted mice for the abundance of CD301b^+^ and CD301b^-^ cDC2 using flow cytometry. Here we found a significant reduction in the absolute numbers of CD301b^+^ cDC2 present in the lungs of *CD11c*^cre^*Crlf2*^flox^, but not of cDC1 or CD301b^-^ cDC2, confirming the TSLP : TSLP receptor axis as a crucial axis for pulmonary CD301b^+^ cDC2 homeostasis (Figure 7C). Taken together our study has identified a pulmonal CD301b^+^ cDC2 niche within the adventitial cuff of the lung to which entry is regulated by PDFGRα^+^SCA1^+^ adventitial fibroblast derived 7α,25-OHC. Within this niche, fibroblastic TSLP supports CD301b^+^ cDC2 survival and *in situ* proliferation (Figure 7D). These data reveal a molecular mechanism where survival of CD301b^+^ cDC2 is a function of subtissular attraction and positioning into a fibroblastic niche within the complex lung microenvironment.

**Figure 7.**
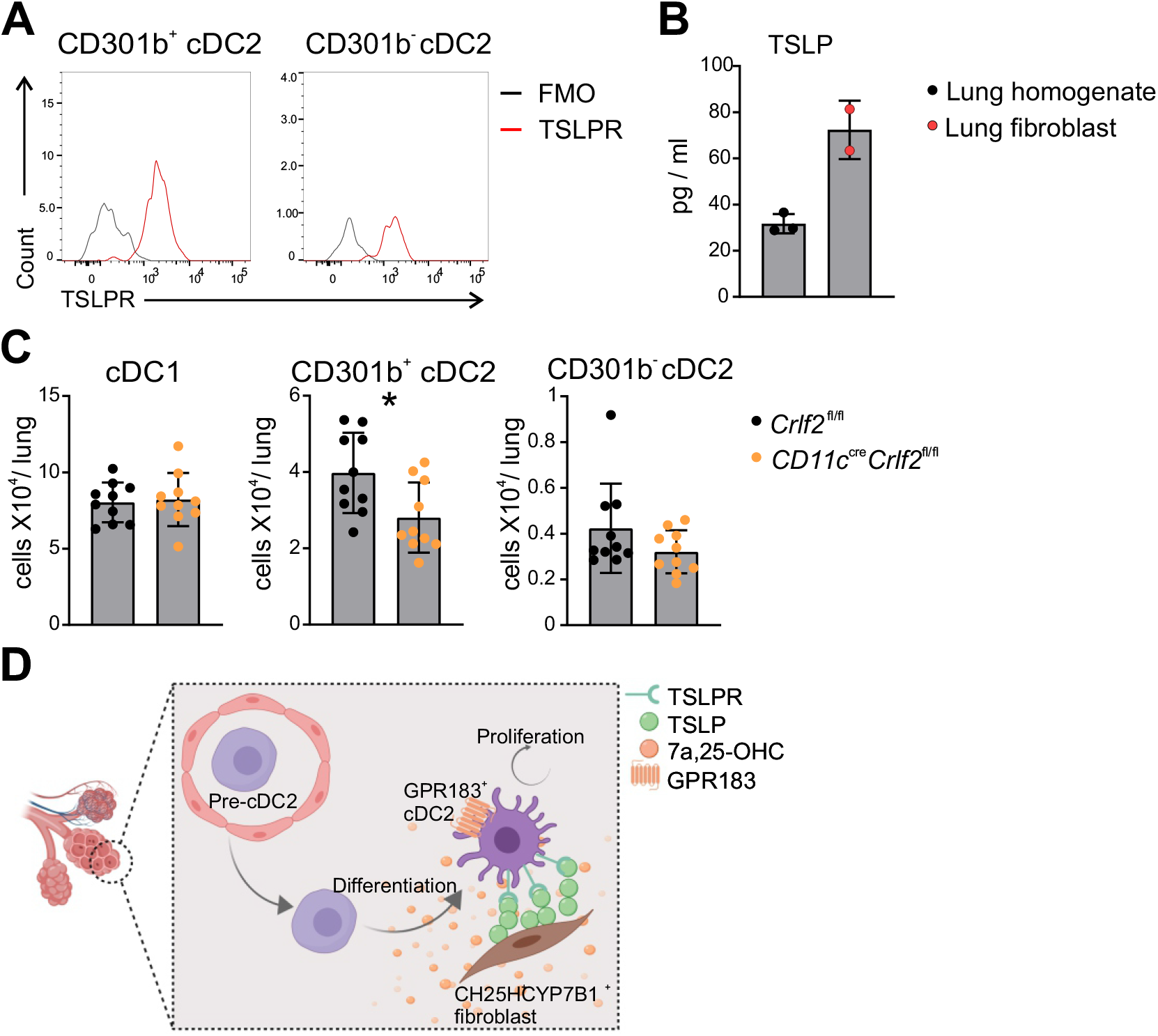
DC specific ablation of TSLP receptor leads to severe reduction of pulmonary cDC2. (A) FACS analysis of TSLPR expression on cDC1, CD301b^-^ and CD301b^+^ cDC2 in WT lung. (B) Analysis of TSLP abundance in lung homogenate (black dots) or within a lung fibroblast culture (red dots) in WT mice. n = 2 or 3. (C) Absolute number of cDC1, CD301b^-^ and CD301b^+^ cDC2 in the lungs of CD11c^cre^*Crlf2*^flox^ (orange) and *Crlf2*^flox^ (black) mice; n = 10. (D) Graphical representation of the proposed mechanism. Each dot refers to one mouse. (A) representative data of two independent experiments. Data in (B) and (C) are pooled from two independent experiments. Data are represented as means ± SD. ∗p < 0.05 by unpaired T-test.

## Discussion

Here we reveal that spatial organization of the pulmonal CD301b^+^ cDC2 network is instructed by 7α,25-OHC – GPR183 chemotaxis. We show that a 7α,25-OHC mediated, GPR183 dependent chemotaxis directs CD301b^+^ cDC2s into a PDFGRα^+^SCA1^+^ adventitial fibroblast niche, which in turn supports CD301b^+^ cDC2 survival via a TSLP receptor dependent interaction. Optimal spatial distribution of cDC subsets is crucial for their function (Cabeza-Cabrerizo *et al*., 2019), however how spatial organization of cDC subsets is achieved remains largely elusive. In the spleen and lymph node GPR183 signalling has been indicated in maintenance and positioning of cDC2s, T- and B-cells (Baptista *et al*., 2019; Gatto *et al*., 2009; Gatto *et al*., 2013; Li *et al*., 2016; Lu *et al*., 2017a). Additionally within lymphoid follicles of the intestine fibroblastic 7α,25-OHC has been shown to attract and organise innate lymphocytes (Emgård *et al*., 2018b). No such effect of GPR183 has been described for non-lymphoid tissue resident cDCs. Despite having no direct pro-survival effect on CD301b^+^ cDC2 GPR183 controls entry of lung resident CD301b^+^ cDC2 to a fibroblastic pro-survival niche providing TSLP to CD301b^+^ cDC2, thereby linking spatial location to survival. In the lung, cDC location mirrors functional specialization, with cDC1 localized close to the epithelial border in order to quickly react to incoming viral pathogens and cDC2s enriched in adventitial areas to survey their environment for extracellular pathogenic cues (Dahlgren *et al*., 2019; Lyons-Cohen et al., 2017). To achieve such integration of functional specialization and microenvironment we identify adventitial fibroblast derived 7α,25-OHC as a subtissular guidance beacon allowing optimal CD301b^+^ cDC2 tissue distribution. In the lung knowledge of stromal derived molecules regulating cDC survival or positioning is scarce. In the spleen and the intestine Notch2 and Ltβr have been reported to be important for cDC2 maintenance, its source being likely of non-hematopoietic origin (Lewis *et al*., 2011; Satpathy *et al*., 2013). More recently retinoic acid and the transition from a mucosal to a epithelial location within the colon was linked to the maturation status of cDC2 (Rivera et al., 2021). Furthermore, in the lung granulocyte macrophage colony stimulating factor (GMCSF) has been associated to the loss of cDC1s and to a lesser extend to that of cDC2 during homeostasis (Greter et al., 2012). More recently it has been demonstrated that an important source of GMCSF in the lung microenvironment are airway epithelial cells, building an intriguing connection between the location of cDC1s within the lung and the production of a cDC1 survival factor, similar to the role of TSLP in regards to CD301b^+^ cDC2 survival (Gschwend et al., 2021).

TSLP has been indicated to be important for type 2 inflammation in the lung, however its homeostatic role has remained unexplored (Connor et al., 2016; Liu et al., 2007). We show that homeostatic *ex vivo* isolated lung fibroblasts produce significant amounts of TSLP and that genetic perturbation of TSLP receptor disturbs CD301b^+^ cDC2 maintenance. CD301b^+^ cDC2 have been shown to be functionally specialized for the induction of Th2 immunity. In the skin interleukin 13 was identified as a crucial mediator of the ability to induce Th2 polarization however cues imprinting those organ-specific functionality within the lung are unidentified (Kim *et al*., 2018; Kumamoto et al., 2013b; Mayer et al., 2021; Sakeen et al., 2015). Therefore TSLP provides an intriguing connection between survival and functional specialization of CD301b^+^ cDC2 in the lung microenvironment.

Fibroblast : macrophage interactions have been well documented in recent years (Franklin, 2021). This paradigm has been conceptualized as the fibroblast : macrophage circuit is important for tissue resident macrophage development, survival and reciprocal functional imprinting. For cDCs such niches have yet to be identified. Importantly evidence suggests that mechanisms exists which confer organ specific cDC functionality, such as retinoic acid signalling, which is important for the induction of tissue-specific anti-tumour T-cell responses in the spleen and the intestine (Klebanoff et al., 2013) or basal interferon alpha signalling in lung and intestine linked to homeostatic and influenza induced functional adaptation of cDC2 and cDC1 during influenza infection (Helft et al., 2012; Schaupp et al., 2020).

Taken together we identify a fibroblast cDC2 circuit, in which niche attraction is regulated by GPR183 dependent migration allowing access to fibroblastic TSLP facilitating CD301b^+^ cDC2 survival. Integration of these and prior results now allows to define a CD301b^+^ cDC2 niche within the lung and establish a framework in which adventitial fibroblasts act as guardians of CD301b^+^ cDC2 survival and maintenance contributing important checkpoints for the initiation of immune responses and the re-establishment of homeostasis.

## Acknowledgments

This study was funded by the Deutsche Forschungsgemeinschaft (DFG, German Research Foundation) under Germany’s Excellence Strategy – EXC2151 – 390873048 (to AS, EK, EM, JH, WK), an Emmy Noether research grant (SCHL2116/1 to AS), DFG research grant (SCHL2116/6-1 to AS). LZ, SS, AS and AP were funded by the Deutsche Forschungsgemeinschaft (DFG, German Research Foundation) – 214362475 / GRK1873/2. E.K. was funded by a fellowship of the Ministry of Innovation, Science and Research of North-Rhine-Westphalia (AZ: 421-8.03.03.02-137069).

---

**Figure S1. CODEX multiplex imaging and flow cytometry to visualize pulmonary stromal and myeloid cell subsets. Related to** Figure 1.

(A) Representative image of the left lobe of a murine lung stained with a 14-plex CODEX antibody panel. Boxed regions indicate the enlarged representative areas in Figure 1A. Scale bar represents 500µm. (B) Representative images of the single stainings of stromal markers used for CODEX. Scale bar represents 100µm. (C) Gating strategy to discriminate various subsets of cDC (cDC1, total cDC2, CD301b^+^ cDC2 and CD301b^−^ cDC2) and other myeloid cell subsets (alveolar macrophages, interstitial macrophages, eosinophils, Ly6C^hi^ monocytes and Ly6C^low^ monocytes). Each subset is indicated with coloured gating. (D) Uniform manifold approximation and projection (UMAP) embedding of pulmonal myeloid cell subsets of a homeostatic WT mouse lung based on (C). Cluster colour indicates classification based on the expression of canonical markers in (C). (E) Representative FACS plots showing GPR183 expression on pulmonary cDC1, CD301b^+^ and CD301b^−^ cDC2 in *Gpr183*^flox-EGFP^ mice.

**Figure S2. Selective reduction of cDC2 in the spleen from *Zbtb46*^cre^*Gpr183*^flox-EGFP^ mice.** Related to Figure 2.

FACS analysis of myeloid cell subsets in lungs of *Zbtb46*^cre^ (black) and *Zbtb46*^cre^*Gpr183*^flox-EGFP^ (blue) mice. Plots show the frequency (A) and cell concentration (B) of myeloid subsets; n = 8-10. FACS analysis of myeloid cell subsets in spleens of *Zbtb46*^cre^ (black) and *Zbtb46*^cre^*Gpr183*^flox-EGFP^ (blue) mice. Plots show the frequency (C) and cell concentration (D) of myeloid subsets in spleen; n = 9-11. Each dot refers to one mouse. Data are pooled from two independent experiments. Data are represented as means ± SD. ∗p < 0.05, ∗∗p < 0.01, ∗∗∗∗p < 0.0001 by unpaired T-test.

**Figure S3. DC lineage specific ablation of GPR183 does not affect cDC development.** Related to Figure 3.

Gating strategy to discriminate MDP, CDP, pre-cDC, pre-cDC1, and pre-cDC2 in the BM (A), blood (B) and lung (C) of WT mice. (D) Frequency of MDP and CDP in the BM of WT (black) and *Gpr183*^-/-^ (red) mice; n = 11 or 12. (E) Frequency of pre-cDC1 and pre-cDC2 in the BM, blood and lung of WT (black dots) and *Gpr183*^-/-^ (red dots) mice; n = 3 or 4. (F) Frequency of total cDC, cDC1 and 2 generated *in vitro* from WT and *Gpr183*^-/-^ BM cultured with Flt3L; n = 9. (G) Frequency of cDC1 and cDC2 generated *in vitro* from WT BM cultured with Flt3L, supplement with or without DMSO, 7α-25-OHC and NIBR189; n = 6. Frequency of Ki67^+^ (H, n = 8) or active caspase 3^+^ (I, n = 4 or 7) pre-cDC1 and pre-cDC2 in the lungs of *Gpr183*^-/-^ (red) and WT (black) mice. Each dot refers to one mouse. Data in (E) are representative data of two independent experiments. Data in (D-H) are pooled from two independent experiments. Data are represented as means ± SD.

**Figure S4. CH25H dependent production of 7α,25-OHC by radiosensitive immune cells does not contribute to maintenance of lung resident cDC2.** Related to Figure 4. (A) Gating strategy for sorting various subsets of stromal cells (epithelial cells, endothelial cells and adventitial fibroblasts) in the WT lung. FACS analysis of lung cDC1, CD301b^-^ and CD301b^+^ cDC2 of CD45.1^+^ bone marrow transplant recipient mice either transplanted with *Ch25h*^-/-^ (pink) or WT (black) bone marrow. Plots show the frequency (B) and cell concentration (C) of cDC1, CD301b^-^ and CD301b^+^ cDC2; n = 6. Each dot refers to one mouse. Data are represented as means ± SD.

**Figure S5. Identification of cDC2 and adventitial fibroblast.** Related to Figure 6.

(A) Expression of *Acta2* and *Myh11* in *GFP*^+^*Col1a1*^+^ cells. (B) Expression of *Pdgfra* in *GFP*^+^*Col1a1*^+^ cells. (C) Expression of *Pi16* and *Gli1* in *GFP*^+^*Col1a1*^+^ cells. (D) Expression of *Ch25h* and *Cyp7b1* in *GFP*^+^*Col1a1*^+^ cells. (E) Expression of *Itgae, Cd86*, *Mgl2* and *Gpr183* in cluster *15*. (F) UMAP embedding of adventitial fibroblast (cluster 9, blue) and cDC2 (cluster 5, red) from untreated murine lung samples generated by Raredon *et al* (GSE133747). (G) NicheNet analysis of potential interactions between receptors on *Mgl2*^+^*Gpr183*^+^ cDC2 and ligands on adventitial fibroblast from untreated murine lung samples generated by Raredon *et al*. (GSE133747).

## STAR METHODS

### RESOURCE AVAILABILITY

#### Lead Contact

Further information and requests for reagents can be addressed to and will be fulfilled by the Lead Contact Andreas Schlitzer.

#### Materials Availability

This study did not generate new unique reagents

#### Data and Code availability

This paper analyses existing, publicly available data. These accession numbers for the datasets are listed in the key resources table. All code used to perform the analysis is available at: https://github.com/JiangyanYu/Zhang_DCs_lung_2021.

### EXPERIMENTAL MODEL AND SUBJECT DETAILS

#### Mice

All animal experiments were approved by Landesamt für Natur, Umwelt und Verbraucherschutz Nordrhein Westfalen. Mice were bred and maintained in a specific-pathogen-free (SPF) condition in Genetic Resources Center (GRC) of the Life & Medical Sciences (LIMES) Institute, University of Bonn, Germany. CD45.1 and *Zbtb46*^cre^ were purchased from The Jackson Laboratory. *Gpr183*^-/-^, *Gpr183*^flox-EGFP^ and *Ch25h*^-/-^ were kindly provided by Prof. Dr. Alexander Pfeifer (University of Bonn). To generate DC conditional depletion of *Gpr183*, *Gpr183*^flox-EGFP^ mice were crossed with *Zbtb46*^cre^ mice. To generate DC conditional depletion of TSLPR, *Crlf2*^flox^ mice were crossed with *CD11c^cre^* mice in an American Association for the Accreditation of Laboratory Animal Care (AAALAC)-accredited animal facility at the Benaroya Research Institute (BRI). Male mice were used in experiments at 8-15 weeks of age. Sex and age-matched littermates were used as wild-type controls.

#### Generation of bone marrow chimeras

Donor BM was extracted from femurs and tibias of mice. BM cells were incubated in RBC Lysis Buffer (Biolegend) for 5 mins at room temperature, then BM cells were resuspended in 1x PBS. 1×10^6^ BM cells in 100 μl of PBS were intravenously injected into lethally irradiated (10 Grey) C57BL/6 recipients within 3 h of irradiation. Recipient mice were rested for 12 weeks before harvesting the lung for assessment of cDC subsets by flow cytometry.

### METHOD DETAILS

#### Cell isolation from tissue

Tissues for flow cytometric analysis were enzymatically dissociated. For dendritic cell analysis, lungs and spleens were cut into small pieces and digested with a 0.2 mg/ml collagenase IV (Sigma-Aldrich) and 0.05 mg/ml Dnase I (Sigma-Aldrich) in HBSS solution for 40 min at 37°C and then mechanically mashed though 70 μm strainers. For pulmonary stromal cell subsets (endothelial cells, epithelial cells and fibroblasts) analysis, lung was inflamed with Dispase II (Merck) and incubated for 10 min on ice in order to dissociate the epithelial layer. The lung was subsequently minced and digested with a 0.2 mg/ml collagenase IV (Sigma) and 0.05 mg/ml Dnase I (Sigma) in HBSS solution for 40 min at 37°C and then mechanically mashed though 70 μm strainers. For lung fibroblasts culture, lungs were cut into small pieces and digested with a 114 U/ml collagenase I (Gibco) and 7.5 U/ml DispaseІІ (Merck) in PBS for 30 min at 37°C with gentle agitation and then mechanically mashed though 70 μm strainers.

#### Lung fibroblast culture

Cells isolated from the WT lung in a way for fibroblast culture were subsequently washed with DMEM supplemented with 10% FCS (Sigma-Aldrich) and penicillin/streptomycin (Gibco) to get crude fibroblasts using “walk-out” protocol. After one day culture of whole lung cells in a dish, unattached were washed away and attached fibroblasts were cultured for 6 more days to get culture supernatant for ELISA.

#### Generation of bone marrow-derived dendritic Cells (BMDCs)

BM cells were harvested from femurs and tibias of 5−8 week-old mice, followed by incubation with RBC Lysis Buffer (Biolegend). BM Cells were counted and resuspend at a density of 1.5 [1 10^6^ cells/ml RPMI medium (PAN Biotec) supplemented with penicillin/streptomycin (Gibco), non-essential amino acids (Sigma), β-mercaptoethanol (Gibco), 10% FCS (Sigma-Aldrich) and 100 ng FLT3L (PeproTech). 4.5 [1 10^6^ cells in the presence or absence of 7α-25-OHC (TOCRIS) or NIBR189 (TOCRIS) were seeded in a 6-well plate per well. DCs were measured by flow cytometry at day7.

#### Flow cytometry

Samples with considerable amounts of red blood cells were treated with RBC Lysis Buffer (Biolegend). Surface staining with respective antibody cocktails was performed in FACS-buffer (0.5% BSA, 2 mM EDTA, 1[1 PBS) and the TruStain FcX antibody (Biolegend) for 30 min at 4°C. Intracellular staining (caspase 3 and Ki67) was performed using the Foxp3/Transcription Factor Staining Set from eBiosciences. AnnexinV was measured according to the manufacturer’s instructions (eBioscience). Live/dead cell discrimination was achieved with Fixable viability dye (1:1000, eBioscience), DRAQ7 (1:1000, BioLegend) or Zombie NIR fixable viability kit (1:1000, BioLegend) for 10min at room temperature. Precision Counting Beads (Biolegend) or AccuCheck counting beads (Invitrogen) were used to determine absolute cell numbers. Details on the reagents used can be found in the Key Resources Table. Stained single cell suspensions were acquired on a LSRII or Symphony A5 flow cytometer (BD Biosciences). Analysis was conducted on FlowJo software.

#### Cell purification

For stromal cell subsets purification, cells from the enzymatically dissociated lung were lysed with RBC Lysis Buffer (Biolegend) before being stained with CD31 (Biolegend), CD45 (Biolegend), Epcam (Biolegend), Sca-1 (Biolegend) and PDGFRα (Biolegend). Cells were labelled with DRAQ7 (1:1000, BioLegend, USA) for 10min at room temperature before suspension in FACS buffer to sort CD45^−^CD31^+^ endothelial cells, CD45^−^Epcam^+^ epithelial cells, CD45^−^PDGFRα^+^Sca-1^+^ adventitial fibroblast. For bulk cDC purification, cells from the enzymatically dissociated lung were stained with CD11c biotin (Biolegend) for 20 min at 4C before labelled with Streptavidin Nanobeads (Biolegend) for 15 min on ice. Cells were then MACS-sorted in antoMACS (Miltenyi) with poss-s selection. Positive fractions containing CD11c positive cells were subsequently stained with Lin (CD3, CD19, Ter119, NK1.1, B220, Ly6G), CD45 (Biolegend), CD64 (Biolegend), SiglecF (Biolegend), MHC2 (Biolegend) and CD11c (Biolegend) for 30 min at 4°C before live/dead labelling. Bulk cDCs were sorted as CD45^+^Lin^−^CD64^−^SiglecF^−^MHC2^+^CD11c^+^. ARIAIII (BD Biosciences, USA) was used for FACS-sorting.

#### Quantitative RT-PCR

RT-PCR was used to confirm transcript abundance differences in FACS-sorted endothelial cells, epithelial cells, fibroblasts and bulk myeloid cells. Total RNA from sorted cells was isolated using the QIAGEN miRNeasy Micro Kit and converted into cDNA via the QuantiTect Reverse Transcription Kit from Qiagen. Gene expression of *Gpr183*, *Ch25h*, *Cyp7b1* and *Hsd3b7* were analyzed using GoTaq qPCR master mix (Promega) and amplification reactions were run on a CFX Connect RT-PCR Detection System (Bio-Rad). Report to the Key Resources Table for complete list of primers used. Expression of each gene was calculated relative to the housekeeping genes PPIA. Relative expression levels (fold changes) were determined by ΔΔCt.

#### Transwell migration assays

FACS-sorted bulk DC suspensions were allowed to transmigrate for overnight across 5 um transwell filters (Corning Costar) toward medium, CCL19 (R&D) or 7a,25-OHC (TOCRIS). 150 μL of 7a,25-OHC (1 mg/mL), CCL19 (200 ng/ml) or media alone with 10% FBS in RPMI were added to the wells of the lower chamber. In co-culture assay, 2×10^4^ PDGFRα^+^ fibroblasts from WT or *Ch25h*^-/-^ mice suspended in 150 μL DMEM medium with 10% FBS were loaded in the wells of lower chamber for one day. 1×10^4^ cells in 70 µL medium were placed into the upper compartment of the transwell. Plates were incubated for overnight at 37°C in 5% CO_2_. Transmigrated cells were enumerated by flow cytometry using counting beads (Biolegend) as reference. The chemotactic index in Fig 1 was calculated via the formula CI = number of cells transmigrated in response to chemotactic factor/number of cells that transmigrated in the absence of chemotactic factor. The chemotactic index in Fig 5 was calculated via the formula CI = number of cells transmigrated in response to chemotactic factor/number of cells that untransmigrated.

#### Confocal lung imaging

Mice were sacrificed by cervical dislocation. The lungs were subsequently inflated with 2% agarose (Promega) via the trachea followed by clamping of the trachea for 5 min. The lungs were then removed and kept on ice-cold PBS for 1 hour, followed by embedding in 4% agarose. Lungs were cut into slices of 200 μm using a vibratome (Leica). Lung slices were blocked, and stained in blocking buffer (Vector Laboratories) for primary antibodies at 4°C for overnight with agitation. Lung slices were stained with anti-PDGFRα (ALY7, 1:500 eBioscience), anti-CD88 (ALY7, 1:500 eBioscience), anti-CD172a (IA4 1:500; Invitrogen) and anti-CD11c (MEC13.3, 1:100, Biolegend). After overnight staining lung slices were washed in PBS for 1 hour followed by Donkey anti-Goat IgG, Alexa Fluor 488 (1:400, Life Technologies) staining at room temperature for 1 hour. Slices were washed and sealed in PBS with nail polish. All preparations were scanned using a Zeiss LSM880 including 405, 488, 561, and 650 laser lines for excitation and imaging with 20X or oil objective. All images were quantified with ImageJ or Imaris.

#### Co-detection by indexing (CODEX) imaging of the murine lung

Mice were euthanized by perfusion in the left ventricle of the heart. Lungs were infiltrated with 50% OCT, excised, and subsequently fixed in 1.3% PFA for 6h at 4°C. Tissue was washed with 1x PBS and dehydrated in 10, 20, and 30% sucrose. After dehydration for 24h, the left lobe of the lung was separated, frozen in OCT, and stored at −80°C. 5µm fresh frozen sections of the left lobe of the lung were stained for CODEX-enabled multiplexed tissue imaging following manufacturer’s instructions. Briefly, coverslips were dried, fixed in ice-cold acetone for 10 min and rehydrated. To reduce the autofluorescence of the tissue, sections were photobleached twice for 1h as indicated in (Du *et al*., 2019) After photobleaching and washing, sections were incubated in staining buffer (Akoya Biosciences), and subsequently stained with a 14-plex CODEX antibody panel overnight at 4°C. After staining, samples were washed twice in staining buffer, fixed in 1.6% PFA for 10 min at RT, and washed three times in 1x PBS. Another fixation step to improve image quality was performed in ice-cold methanol for 5 min, followed by three washing steps with 1x PBS. Finally, samples were fixed with BS3 crosslinker (Sigma Aldrich) for 20 min at RT to preserve antibody binding during imaging and washed as in the previous fixation steps. Specimens were stored in CODEX storage buffer (Akoya Biosciences) at 4°C for up to 1 week or directly imaged. Prior to imaging, fluorescently labeled reporters were diluted and pipetted into a black 96-well plate, following manufacturer’s instructions (Akoya Biosciences). Image acquisition was performed with a Zeiss Axio Observer widefield microscope (Carl Zeiss) using a 20x air objective (NA 0.85). A multicycle acquisition experiment was set using the CODEX Instrument Manager (CIM, Akoya Biosciences) following the manufacturer’s instructions (Akoya Biosciences). Multi-channel images of several tiles with a z-spacing of 1.5 µm were acquired, covering the entire lung section.

#### CODEX image processing and analysis

Images were exported using the CIM data transfer tool (Akoya Biosciences). Image processing was done using the CODEX Processor (Akoya Biosciences). Briefly, images were stitched, optical shading was corrected, background (calculated using blank cycles) was subtracted, and deconvolution with Microvolution was performed. Finally, a single best focus image was generated for further analyses. Nuclear segmentation was performed using HALO AI (Indica Labs, Albuquerque, NM, USA). DAPI counterstain was selected as a reference for this purpose. Additionally, EpCAM staining was used to improve the definition of cell boundaries at the aerial epithelium. After training, the HighPlex FL module of HALO (Indica Labs) was employed to define the threshold values for each of the antibodies of the panel. These values were further used to classify the cells into phenotypes. cDC1 were defined as CD45^+^F4/80^-^MHCII^+^CD11c^+^CD11b^-^Xcr1^+^ objects, whereas cDC2 were defined as CD45^+^F4/80^-^MHCII+CD11c^+^CD11b^+^CD172α^+^ objects. cDC2 subpopulations were further separated based on the threshold values defined for CD301b staining. Layers delineating bronchi, bronchioles, alveolar spaces and adventitial cuffs were manually annotated. Absolute cell numbers for each of the annotated layers were calculated and normalized by the area.

#### Single cell transcriptomic Receptor : Ligand analysis

To identify a molecular crosstalk between cDC2 and adventitial fibroblast in silico, we made use of two recently published ((Tsukui *et al*., 2020), GSE132771; (Raredon *et al*., 2019), GSE133747). First Gpr183^+^ cDC2 and adventitial fibroblasts were identified from this dataset. In brief, the downloaded matrix (GSM3891616-GSM3891619) was processed using Seurat (V4.0.1) (Hao *et al*., Cell, 2021). As described in Tsukui *et al*., cells with fewer than 250 detected genes or cells with >10% of mitochondrial gene counts were excluded in the quality control step. Graph-based clustering was performed using the first 20 dimensions and a resolution setting of 0.6. In order to identify *Gpr183^+^* cDC2s in this dataset, we first used cDC marker gene, *Cd86* and *Itgae*, to identify cDCs (cluster 15). Within the DC cluster, we defined cells that express *Mgl2* and *Gpr183* as *Gpr183*^+^ cDC2. Next to identify adventitial fibroblasts, we extracted GFP^+^ cells as described in Tsukui *et al*. paper and performed graph-based clustering analysis on this subset of cells using Seurat. The adventitial fibroblasts were defined using the marker genes *Pi16* and *Pdgfr*α. Next, we predicted the cellular interaction between *Ch25h*^+^*Cyp7b1*^+^ adventitial fibroblasts and *Gpr183*^+^ cDC2 using NicheNet (V1.0) that links ligands from sender cells to target genes in receiver cells (Browaeys *et al*., 2020). As the target gene set, we considered the list of genes that are differentially expressed between *Gpr183*^+^ cDC2 and all other cells from the two untreated lung samples isolated from GSE132771 (Wilcoxon rank-sum test; adjusted P[≤[0.05 and log2 fold change[≥[1). The top 30 ligands that were expressed in adventitial fibroblasts and were with the highest Pearson correlation to the target genes were included for downstream analysis. The ligand-receptor interaction was visualized in a circos plot from the circlize package (V0.4.12) according to the instructions from NicheNet. To further validate our findings, we performed in silico analysis of cellular crosstalk between cDC2 and adventitial fibroblasts using another recent published dataset (GSE133747). From this dataset we utilized samples GSM3926539 and GSM3926540, and imported them into the Seurat pipeline. Receiver cells were identified using expression of *Itgae*, *Cd86* and *Mgl2*, while sender cells were extracted via marker genes *Pdgfr*α and *Pi16*. Simultaneously, the cellular interaction modelling was performed as described above using NicheNet.

### QUANTIFICATION AND STATISTICAL ANALYSIS

Statistical analysis was performed using Prism software (GraphPad). Statistical significance was determined using the two-tailed unpaired Student’s T-test with at least 95% confidence. One-way analysis of variance was initially performed to determine whether an overall statistically significant change existed before using unpaired T-test. Data were shown as mean ± SD. Statistical significance was considered for *p < 0.05, **p < 0.01, ***p < 0.001, and ****p < 0.0001.

## ADDITIONAL RESOURCES

### Author contributions

Conceptualization: LZ, AS; Formal Analysis and Investigation: LZ, SS, SS, BD, VM, TQ, IM, SU, BC, SR; Writing: LZ, JY, AS; Supervision: AS, EM, JH, AP, EK, WK, SZ. All authors contributed to the manuscript.

### Declaration of Interests

All other authors declare no financial interest.

## Notes

### Competing Interest Statement

The authors have declared no competing interest.

