## Supplementary figures and images for "GPR183 targets lung-resident CD301b^+^ conventional dendritic cells type 2 to a subtissular TSLP – TSLP receptor mediated survival niche within the adventital cuff"

### Supplemental Figure 1

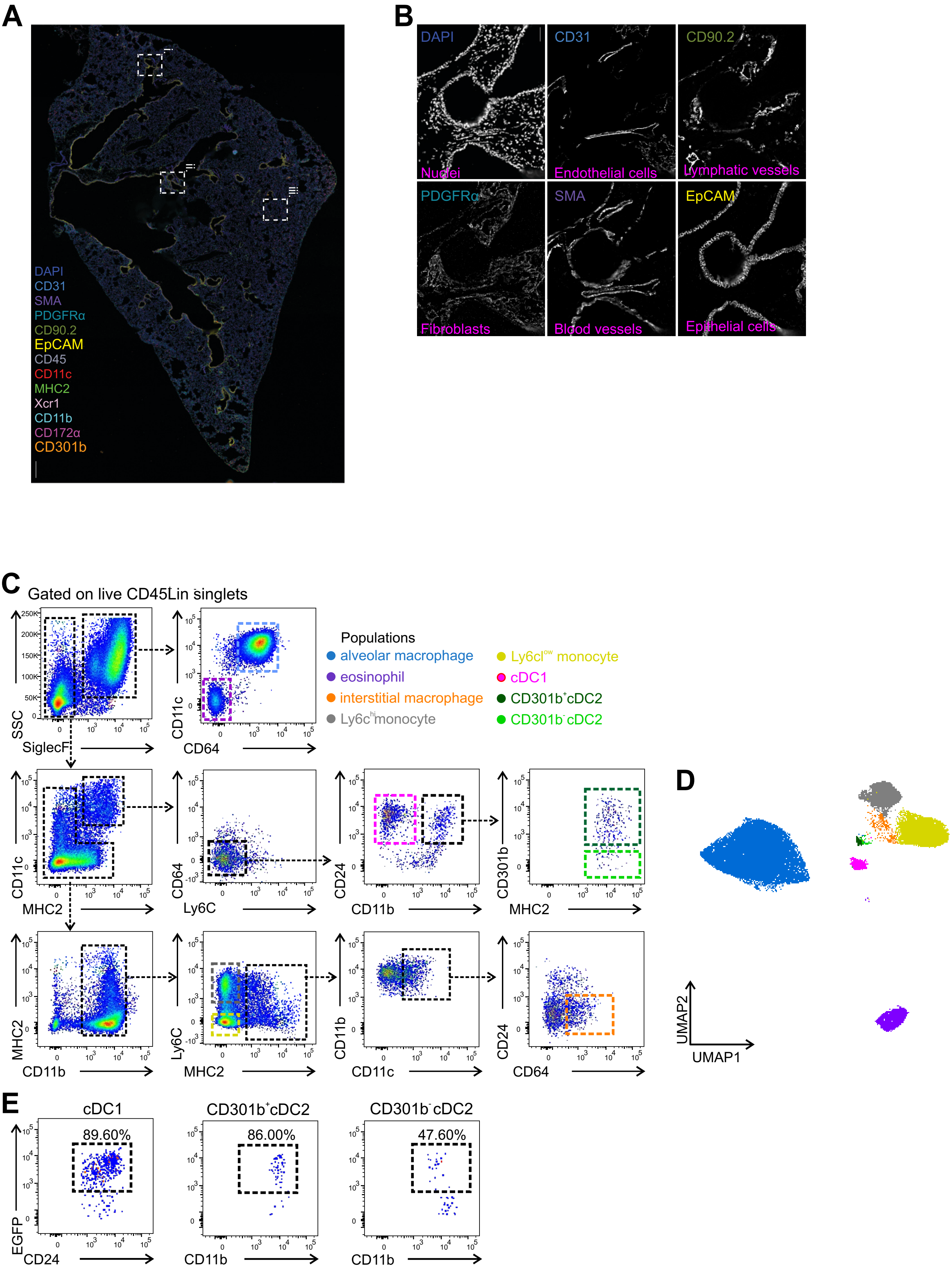

### Supplemental Figure 2

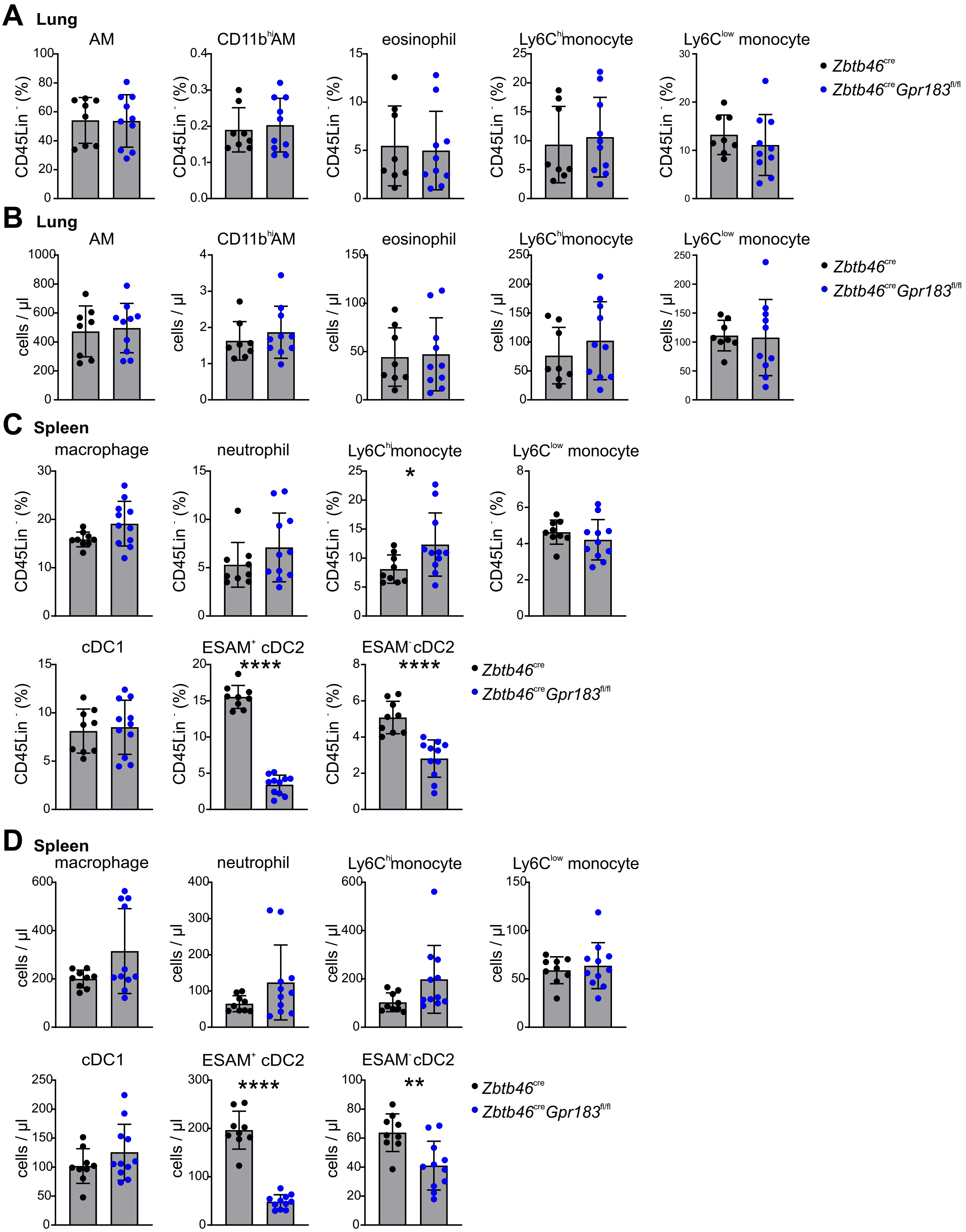

### Supplemental Figure 3

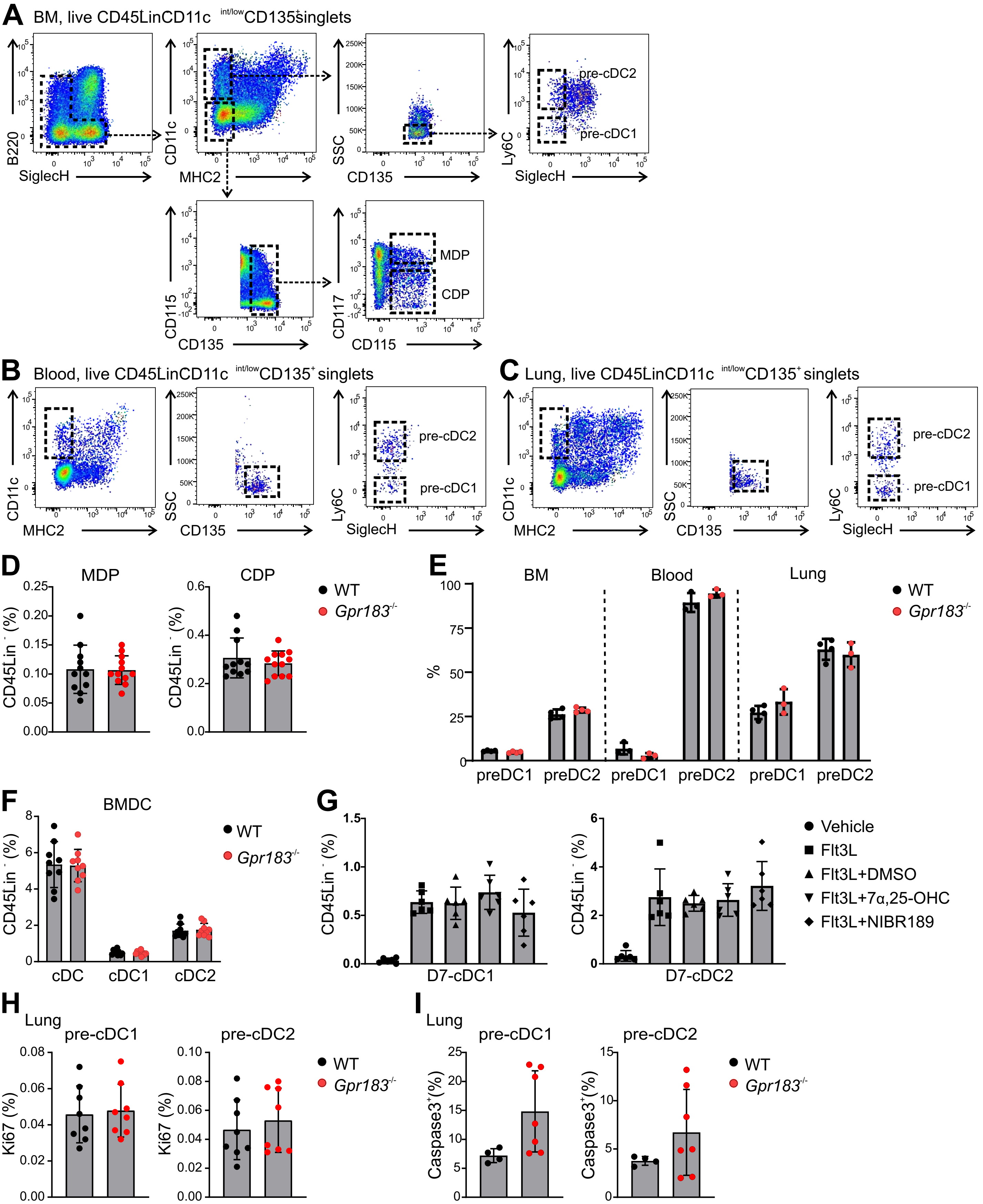

### Supplemental Figure 4

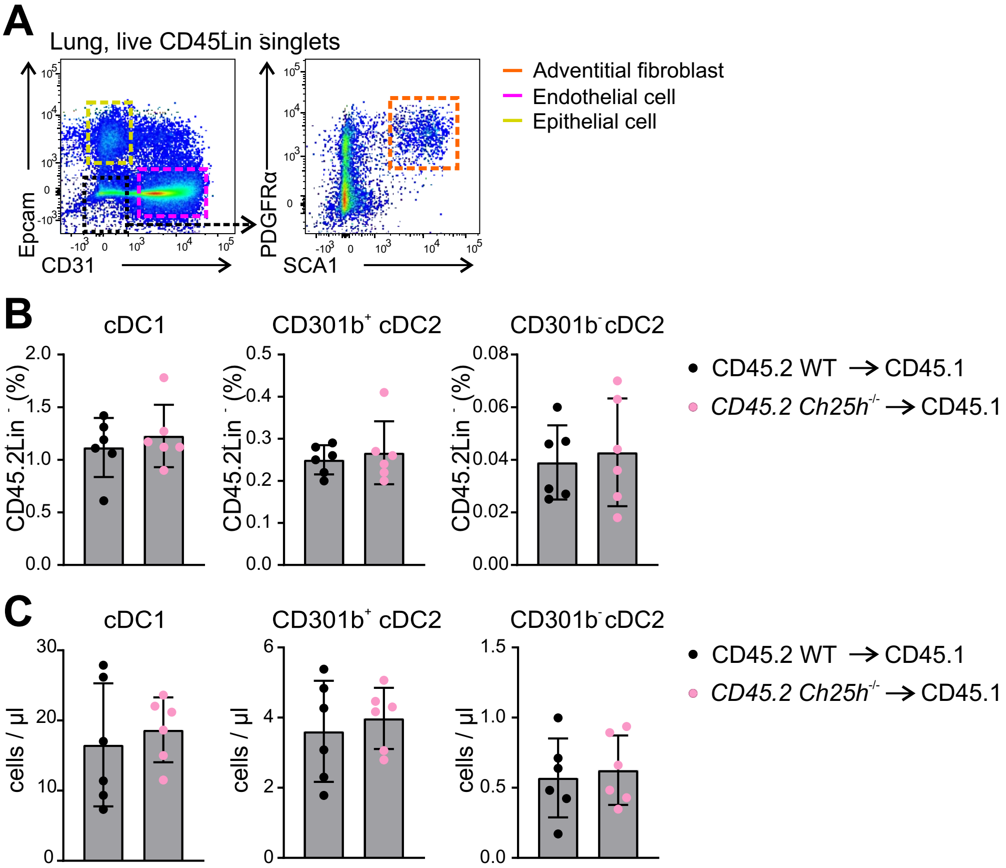

### Supplemental Figure 5

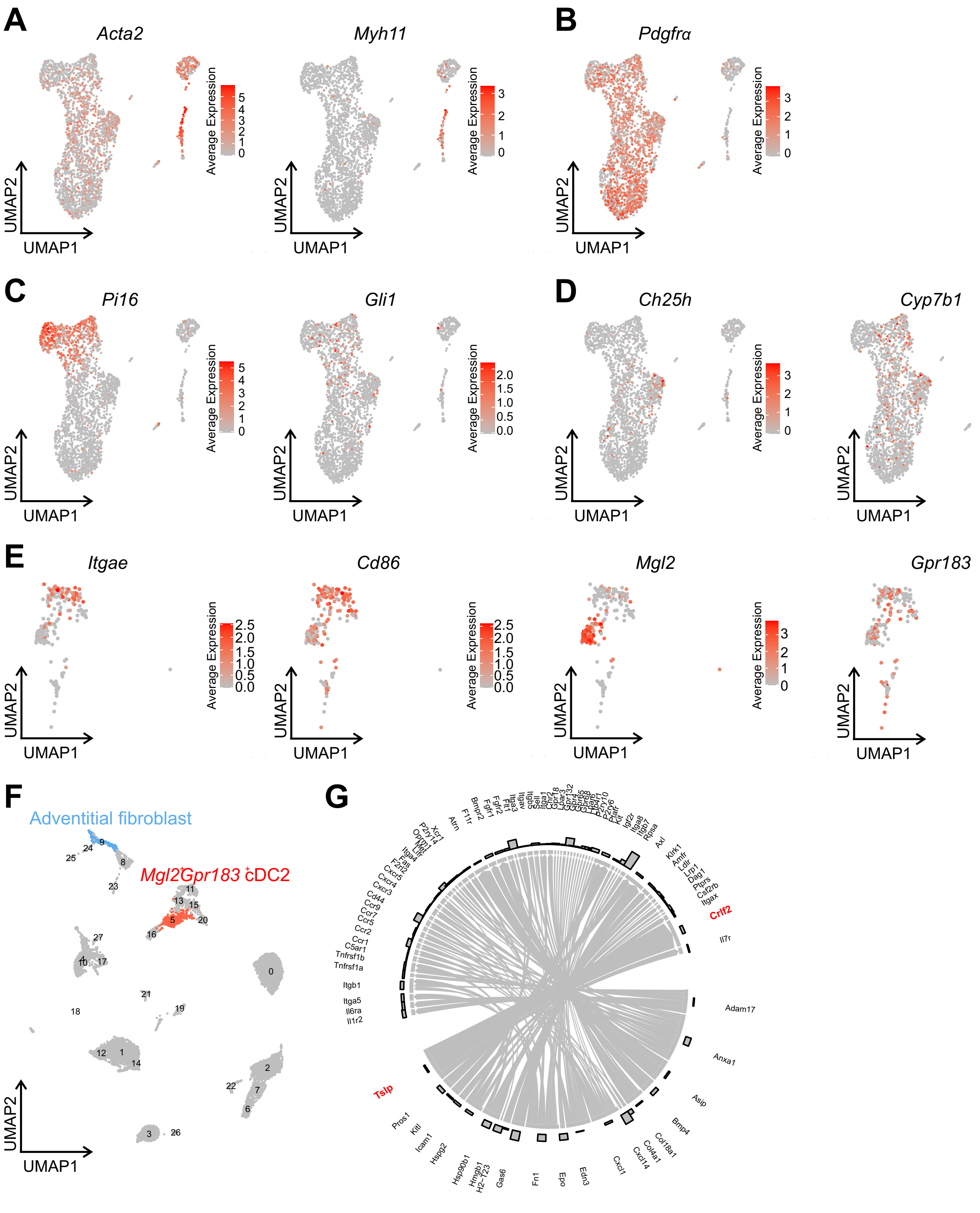
